# The sense of agency from active causal inference

**DOI:** 10.1101/2024.01.29.577723

**Authors:** Acer Yu-Chan Chang, Hiroki Oi, Takaki Maeda, Wen Wen

## Abstract

This study investigates the active component of the sense of agency (SoA), positing that SoA is fundamentally an outcome of active causal inference regarding one’s own actions and their impact on the environment. Participants controlled visual objects via a computer mouse, with tasks designed to test their ability to judge control or detect controlled objects under varying noise conditions. Our findings reveal that participants formed high-level, low-dimensional action plans that were idiosyncratic across but consistent within individuals to infer their degree of control. Employing transformer-LSTM-based autoencoders, we captured these action plans and demonstrated that the geometrical and dynamical properties of these action plans could predict behavioural profiles in the tasks with remarkable accuracy. This suggests that participants’ sense of control is shaped by actively altering action plans, viewed as generating causal evidence through intervention. Further, participants proactively expanded the diversity of their action plans, facilitating the exploration of available action plan options while accumulating causal evidence for the inference process. Contrarily, patients with schizophrenia exhibited reduced action plan diversity, suggesting impaired active control inference and detection of self-relevant cues. These findings offer a more comprehensive understanding of the sense of agency, deeply rooted in the process of active causal inference.

## 1 Introduction

Recent research suggests that various aspects of conscious experience are associated with the outcomes of internal inferential processes. The predictive processing theory of consciousness, for instance, posits that our conscious experiences are essentially the ’best guesses’ or the most probable inferences drawn from our internal models about the external world [Hohwy, 2013, Seth and Hohwy, 2020].

This notion can be extended to the sense of agency (SoA)—the conscious feeling of control over one’s actions and their outcomes [Gallagher, 2000]. We constantly infer our causal role in bodily and environmental changes, thereby constructing an estimated understanding of our influence over these changes that results in the SoA. This notion well aligns with the Bayesian inference framework of SoA as recently proposed in recent literature [Yano et al., 2020, Legaspi and Toyoizumi, 2019, Moore and Haggard, 2008, Wolpe et al., 2013, Moore and Fletcher, 2012, Moore, 2016, Wen and Imamizu, 2022, Lush et al., 2019, Synofzik et al., 2009]. In essence, causal inferences are fundamental to our comprehension of ourselves as agents adapting to and interacting with the environment.

However, distinct from perceptual inference, the inference associated with the sense of agency necessitates action execution. This differs from the straightforward comparison of predictive and actual sensory feedback as proposed by the classical comparator model [Frith et al., 2000, Blakemore et al., 1999, 2002]. The incorporation of action can potentially benefit the inferential process by allowing for the deliberate choice of action policies. These policies can promote the generation and collection of causal evidence and allow exploration of controllable actions within the action space to optimise inferential outcomes. Although it is of critical importance, the literature still lacks a thorough dissection of the action policies that are central to the sense of agency.

The current study examines the emergence of the sense of agency as a product of active causal inference regarding our controllability over the environment. Specifically, we hypothesise that action plays an essential role in inferring one’s causal role, marking a departure from the passive nature of perceptual causal inference. This notion of actively utilising actions for inference is agreement with the notion of active inference in the framework of the free energy principle [Friston et al., 2009, Friston, 2010b, Friston et al., 2013].

The hypothesis underpinning our study posits that: 1. The capacity to actively formulate action plans enables the selection of action policies less susceptible to environmental noise, thus enhancing the reliability of our inferences; 2. Through action, we can transcend mere causal inference from correlation by generating causal evidence via **intervention** to strengthen the inference of control capacity; 3. Action facilitates an increase in the diversity of executable actions, which, in turn, allows for both a broader exploration of the action space and the accumulation of more causal evidence to support inference.

To test these hypotheses, we employed data collected in one prior study [Oi et al., 2023] encompassed two tasks crafted to investigate the SoA [Wen et al., 2020a, Wen and Haggard, 2018, 2020, Wen et al., 2020b]. A pivotal aspect of our dataset is the inclusion of data from both healthy individuals and patients with schizophrenia—a demographic known for impaired agency judgment [Haggard, 2017, Frith et al., 2000]. This allows for a comprehensive assessment of how varying levels of the SoA, due to schizophrenia, affect the capacity to actively engage in causal inference.

Both task involved participants freely moving a computer mouse for a period of 5 seconds. The first task, the *control detection task*, asked participants to identify the target dot from three dots on the screen, including two distractors (Figure 1a). The second task, the *control judgment task*, required participants to indicate whether they felt they were able to control the motion of a target dot (Figure 1b). In order to manipulate the level of control, two types of spatial control noise were introduced: degree of control (30%, 55%, and 80%) and affine transformations (i.e., rotations of 0 or 90 degrees) were applied to the movement direction of the target dot (Figure 1c). In this article, we use the term *volitional* and *displayed* to refer to the data obtained from participants’ unaltered movements and the one that had undergone experimental noise and was subsequently presented on the screen, respectively (see 5.3.2).

**Figure 1:**
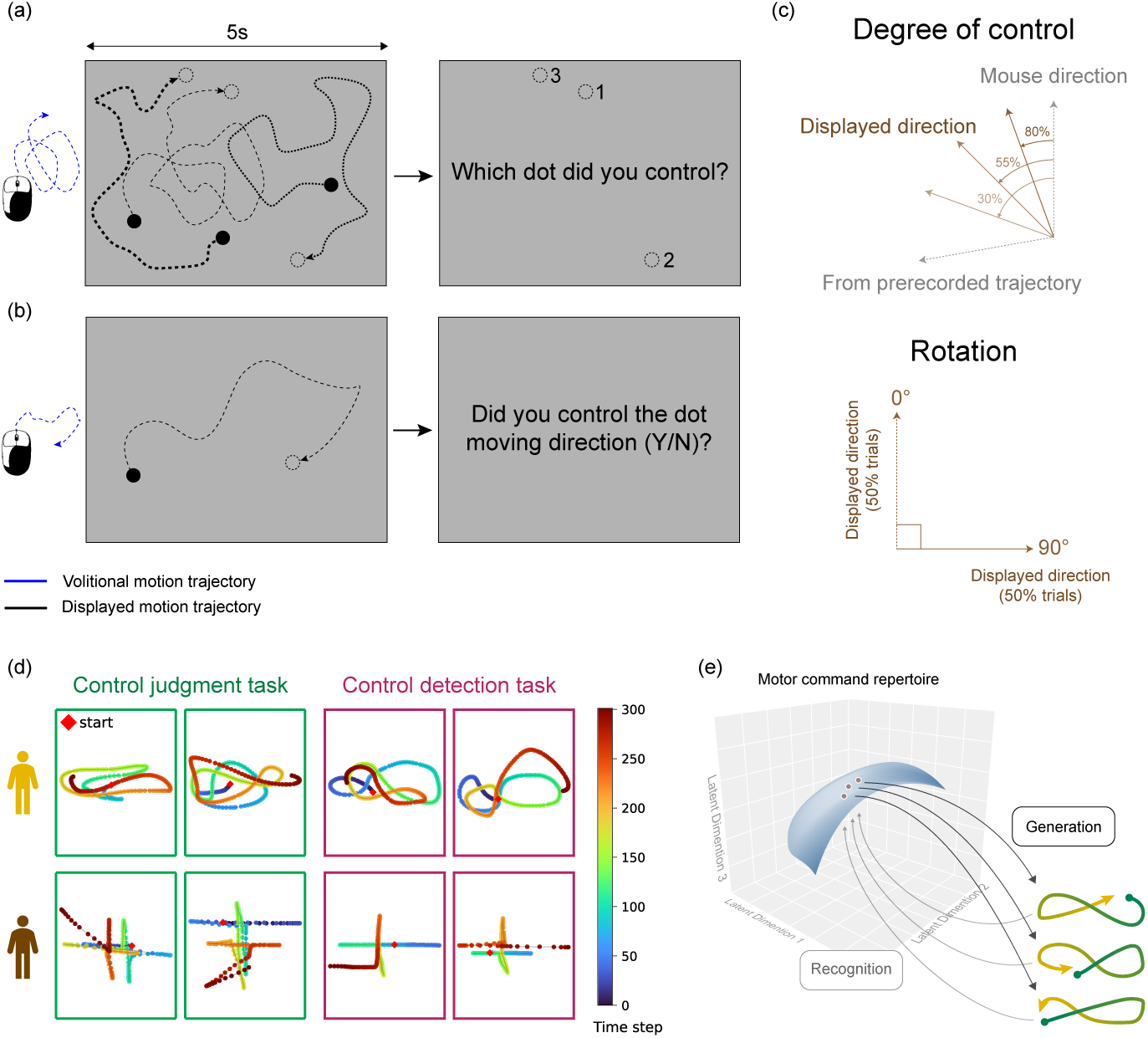
Illustration of task designs and participants’ pattern-creation behaviours. (a) The control detection task. Participants moved the mouse to partially control the motion of the target dot among two distractor dots with predetermined paths. After 5 seconds, participants selected the target dot. (b) the control judgment task. Participants moved the mouse for 5 seconds to control the motion of the target dot, then judged whether they felt they could control the dot’s movement. “Volitional” refer to the data obtained from participants’ unaltered movements and “displayed” to refer to the one that had undergone experimental noise and was subsequently presented on the screen. (c) Two types of interferences applied to the motion direction of a target dot: degree of control and rotation. The degree of control varied between 30%, 55%, or 80% and was determined by the proportion of the participant’s mouse movement that influenced the dot’s moving angle. The 90° rotation op motion direction was adding to 50% of the trials. The two interference factors result in six conditions. (d) The volitional motion trajectories generated by two exemplary participants exhibiting pattern-creation behaviours. Each panel depicts the motion trajectory of a single trial. The examples highlight the idiosyncrasy and consistency of participants’ patterns across trials and tasks. (e) A conceptual diagram depicting the recruitment of similar trajectory patterns suggesting that the actions originate from a low-dimensional manifold of motor command repertoires

In accordance with our hypotheses, we first asked whether the participants actively perform specific **action plans**, instead of random movement, to fight the local noise we applied to the motion directions of the target dot. We then asked whether the participants actively use the action plan to generate causal evidence through intervention, that is, by actively changing their action plans. We predicted that participants would monitor the corresponding dynamic changes in the environment, which supports the inference of control capacity. Lastly, we asked whether the participants would actively increase the diversity of action plans to gather and explorer more causal evidence for the inference.

The study employs a transformer-LSTM-based autoencoder [Hinton and Salakhutdinov, 2006, Vaswani et al., 2017, Greff et al., 2016] to capture and quantify the individual’s idiosyncratic action plans. The transformer-based encoder is well-suited for capturing long-term dependencies, enabling it to encode motor sequences and recognize action patterns. The decoder, constructed with LSTM (Long Short-Term Memory) layers, acts as a generative model that produces the motor sequence based on high-level (in contrast to moment-to-moment motor commands) and abstract (representing overall patterns of movement) motor commands, as well as the current state. By transforming long motor sequences into low-dimensional action plans, our approach allows us to analyse and understand the sense of agency at the abstract action plan level.

Finally, our study incorporated a comparative analysis between healthy individuals and schizophrenia patients [Maeda et al., 2012, 2013], revealing novel insights into how schizophrenia impacts the sense of agency. This comparison, particularly focusing on active causal inference in patients, highlights a previously under-explored area in research. Our findings emphasize the importance of active causal inference in comprehending the sense of agency, especially within the context of psychiatric conditions.

## 2 Results

### 2.1 Participants exhibited pattern-creation behaviours

Through qualitative observation, we found that participants manifested divergent pattern-creation behaviour in both the control detection task and the control judgment task. The patterns generated by each participant were found to be consistent across trials and both tasks as illustrated in Figure 1d. This intriguing behaviour also suggests that a high-level but low-dimensional motor command repertoire is employed in the motor generation process, (Figure 1e). The initial qualitative findings indicate that participants may create the macro-level action plans to enhance sensory signals caused by self-generated action while mitigating local noise.

### 2.2 Action plan analysis

To capture the motor command repertoire, for each individual, we trained a transformer-LSTM based autoencoder (Figure 2a, more details see Figure 7). The autoencoder was trained on the volitional motor sequences recorded from the control judgment task. Volitional motor sequences are those intrinsically generated by participants reflecting their intentional movements before the introduction of any experimental noise (Figure 2d). The low-dimensional bottleneck (hidden) layer of the autoencoder was utilised to extract high-level and low-dimensional motor commands, which we termed the *action plan* (Figure 2e). This analysis aims to delineate the contribution of these action plans to the participants’ subjective experience of agency.

**Figure 2:**
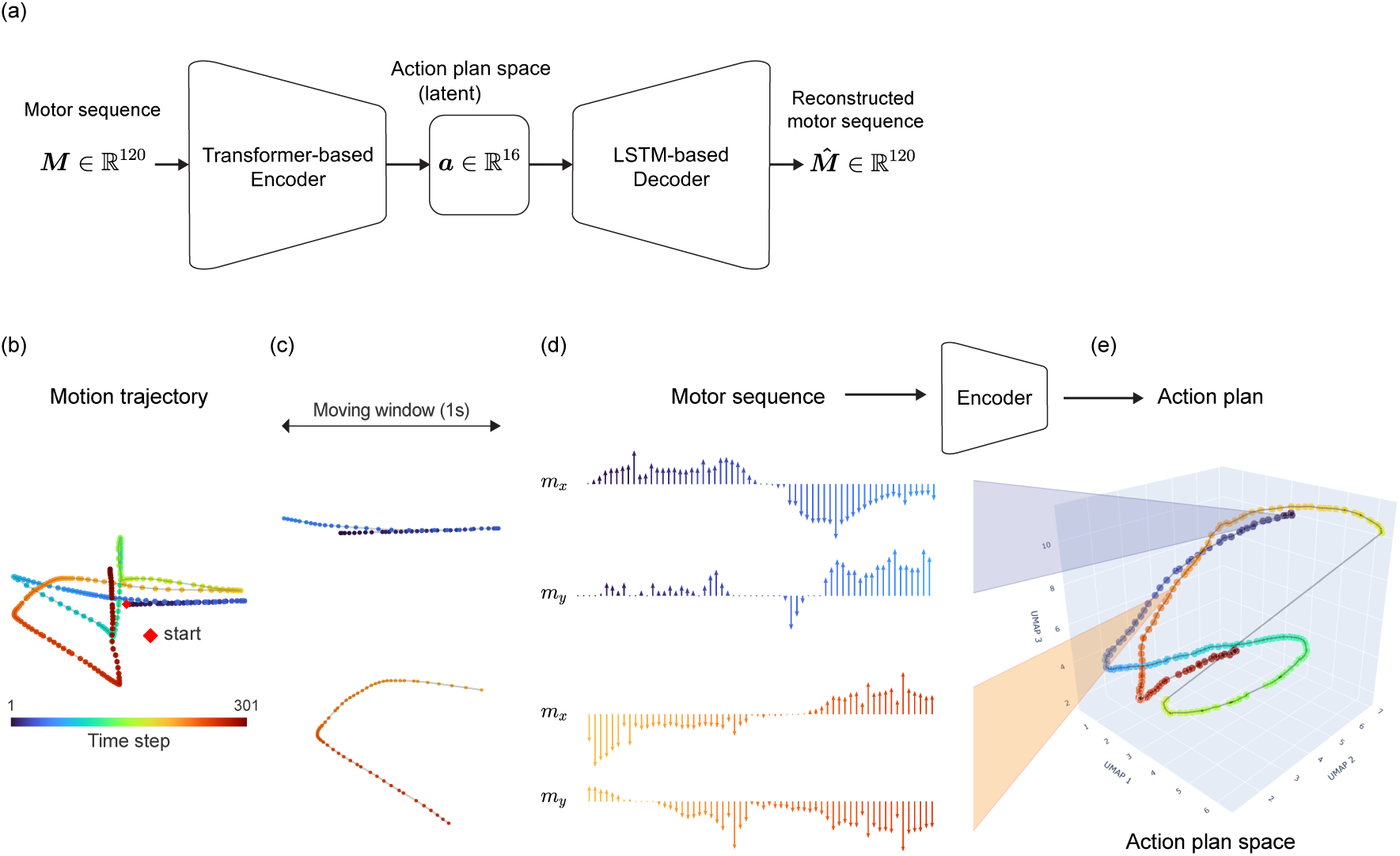
Transformer-LSTM-based autoencoder and data processing pipeline. (a) The autoencoder consists of a transformer-based encoder and an LSTM-based decoder. The encoder transforms the 120 dimensional motor sequence ***M*** into a low-dimensional (16) action plan ***a***, while the decoder reconstructs the motor sequence ***M̂*** from the action plan. (b) A motion trajectory from a representative trial. (c) The trajectory is sliced into subsequences using a moving window with 1-second window size and 1-sample step size. This figure illustrates two example subsequences. (d) The x and y directions of the motor sequences of the two example subsequences. (e) Mapping the motor sequences to action plans with the encoder. Note that the Uniform Manifold Approximation and Projection (UMAP, McInnes et al. [2018]) here for dimension reduction is only used for the visualisation purposes and not involved in data processing.

Therefore, by utilizing the trained encoder, we were able to quantify the dissimilarity between action plans by computing the (Euclidean) distance between them (Figure 3a-b). If this pattern-creation behaviour is indeed a product of the participants’ active attempt to infer their capacity to control, we would anticipate the geometric relationship could be critical to the underlining inference process regarding participants’ agency perception. As such, in our subsequent analyses, we utilised the encoder to map both volitional and displayed (see 5.3.2) motor sequences to action plans and conducted the following analyses within the action plan space.

**Figure 3:**
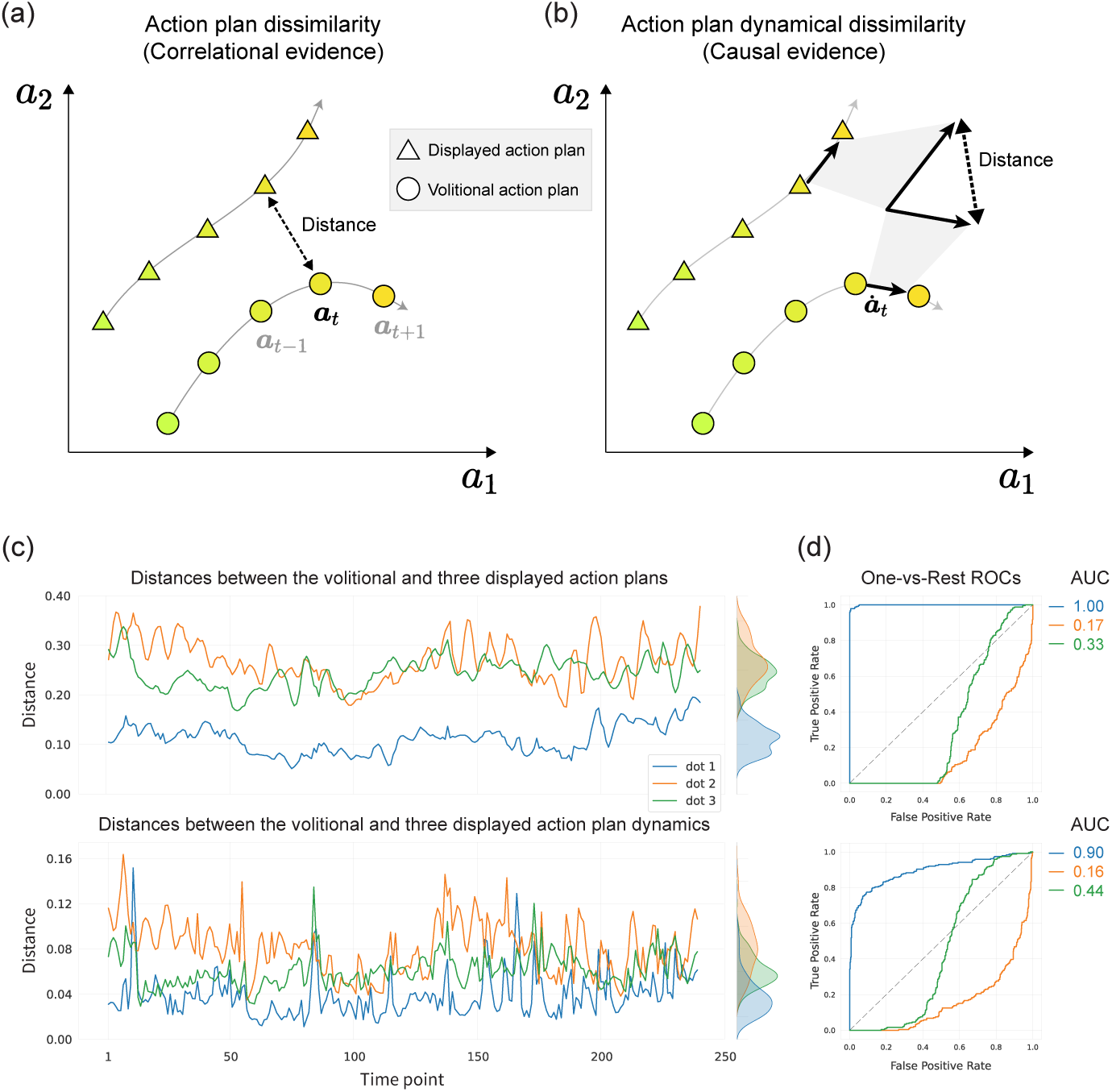
Quantitative Analysis of Action Plan (a) At any given time point *t*, the dissimilarity between action plans can be measured by calculating the Euclidean distance between the volitional action plan and the displayed action plan. This measurement provides correlational evidence to support active causal inference processes. (b) The dynamical dissimilarity between action plans can also be quantified by the geometric distance between the volitional and displayed action plan dynamics, serving as causal evidence for the inference. (c) The upper panel shows the Euclidean distances between the volitional and three displayed action plans from an example trial in the control detection task. The lower panel shows the distances between the volitional and displayed action plan dynamics. (d) The Receiver Operating Characteristic (ROC) curves for one-versus-rest classifications are shown, with the corresponding Area Under the Curve (AUC) values illustrating the discriminative power of the distance measures. In this example, dot 1 was selected as the target due to the highest AUC within the six.

### 2.3 Results from Healthy Participant Group

We commenced our analysis of the results by first focusing on the healthy participant group. Following the detailed analysis of the healthy participant group, we extended our investigation to include the patient group.

#### 2.3.1 Geometric Relationships in Action Plans as Predictors of the Sense of Agency in Control Detection

To determine whether the autoencoder we trained effectively captures unique high-level motor commands exhibited by participants across tasks, we first examined whether the geometric relationship between action plans can predict participants’ performance in the control detection task. It is worth noting that the autoencoders were trained using data from the control judgment task, and therefore any effects observed in the control detection task would indicate the successful generalisation of the trained autoencoders.

We predicted that if participants inferred their capacity for control at the action plan level, they would consider the distances between their volitional and displayed action plans (Figure 3a). We also posited that participants would not only rely on correlational evidence, i.e., the similarity of action plans, but also causal evidence, i.e., the dynamics (changes) of action plans (Figure 3b). Specifically, they would monitor whether changing their action plans could lead to corresponding changes in their environment in the same direction, i.e., the effect of intervention. We define *action plan dynamics* as the difference between temporally adjacent action plans. By considering both correlational and causal evidence, individuals could make a more precise causal inference, which would lead to better detection of self-relevant signals.

In each trial, a one-second sliding window was applied to the volitional and the three displayed motor sequences to obtain a series of one-second subsequences (Figure 2c, and see 5.5.1 for details). The action plan of each subsequence was computed by running the encoder on the subsequence. At each sample time point of the action plan sequence, we calculated the distances between the volitional action plan and the displayed action plans. This results in three distance series. Similarly, the computation was applied to the action plan dynamics (Figure 3c).

To predict participants’ response in each trial of the control detection task, we employed the one-vs-rest receiver operating characteristic (ROC) curves to quantify the separability of three distance series Hanley and McNeil [1982]. The areas under curve (AUC) were then calculated for each ROC curve to provide a scalar measure of each distance series. We repeated this computation procedure for both the action plan and action plan dynamics, resulting in six AUC values in total. The distance series with the highest AUC among the six was selected as the most salient self-relevant signal, enabling us to predict the participants’ response in each trial (Figure 3d and see 5.6 for more details).

As the inferred decision-making was modelled on a trial-by-trial basis, it allowed us to predict accuracy for each participant in each condition. We hypothesised that if the participants utilised the action plan to actively infer their control capacity and identify the target, and if this use of the action plan was a common computational principle, the predicted accuracy would positively correlate with the actual accuracy across participants and all conditions.

The results (Figure 4a,d) show a highly significant correlation between predicted accuracy and ground truth accuracy for the healthy group (*r* = 0.901*, p <* 10*^−^*^54^). The high correlation between the predicted accuracy and the ground true accuracy for the healthy group suggests that the geometric relationship between action plans in the action plan space can well explain responses across participants and conditions. Notably, this finding also suggests that the two types of noise applied in the study can be well explained by a single concise principle.

**Figure 4:**
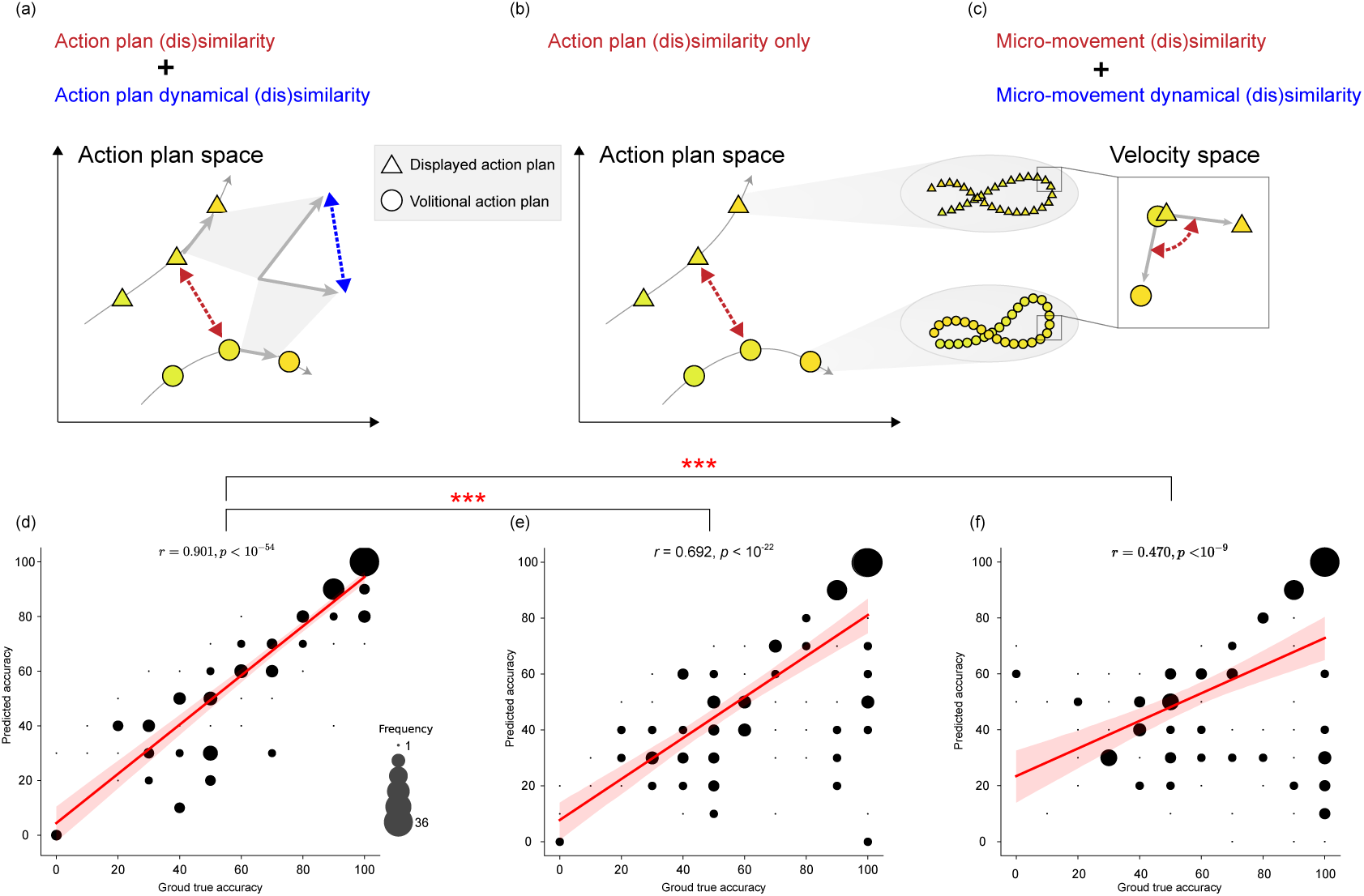
Predicted versus actual task performance in the control detection task for the health group. (a)(d) Demonstrates a robust positive correlation between the predicted and actual accuracy among healthy participants across participants and conditions in the control detection task, signifying that action plans together with action plan dynamics can predict responses effectively. (b)(e) Demonstrates a discernible, yet notably diminished correlation between the predictive accuracy, which excludes action plan dynamics, and the actual task performance, underscoring the significant influence of action plan dynamics in accurate prediction. (c)(f)The correlation indicating a significant but lower predictive accuracy based on micro-movements when compared to action plans suggesting that action plans play a more prominent role in shaping the sense of agency than micro-movements in the control detection task.

Upon a qualitative examination of the aggregated behavioural profile at the group level (Figure S2a), it becomes evident that the predicted accuracy not only reflects the actual accuracy but also effectively mirrors the impacts of the two noise types on the inference process and conscious experience of control.

#### 2.3.2 Assessing the Impact of Causal Evidence and Intervention Through Action Plan Dynamics in the Control Detection Task

We then examined whether the action plan dynamics played a significant role in the control detection task. We predicted that if participants actively inferred their causal role in environmental dynamics, they may change the action plan to exert intervention and then monitor its consequence. To test this hypothesis, we performed the same analysis as above, except only using the information of action plan instead of information from both the action plan and the action plan dynamics(Figure 4b).

The results revealed a notable decrease in the correlation between predicted accuracy and actual accuracy among the healthy group when solely considering action plan information (*r* = 0.692*, p <* 10*^−^*^22^, 4e). This significant decline (*z* = 5.36*, p <* 10*^−^*^7^), underscores the importance of action plan dynamics in the participants’ capacity to detect self-relevant signal during the control detection task.

It is important to highlight that action plan dynamics are derived from a rolling window approach, where each action plan is computed with a 1-sample step difference from the previous one. Although these differences may appear minor in motor sequences, they become substantially influential at the action plan level, markedly impacting self-relevant information detection. This again demonstrates the importance of causal and interventional evidence in the underlying active inference process of the sense of agency.

In sum, the analysis of the action plans in the control detection task yielded results that suggest that participants’ sense of agency can be accurately explained by the geometric relationships within the action plan space, even under varying conditions. These findings provide strong support for our hypothesis that the active creation of action plans plays a crucial role in helping participants infer their control capacity over the environment. Furthermore, the observed influence of action plan dynamics on the sense of agency reinforces our hypothesis that participants took into account both correlational and causal evidence when forming their sense of agency.

In addition, the result demonstrates the autoencoder’s generalizability: trained on control judgment task data, it effectively captures action plans and predicts detection accuracy in the control detection task. This indicates both methodological robustness and consistency in participants’ action plans across tasks.

#### 2.3.3 Geometric Relationships in Action Plans as Predictors of Agency Sense in Control Judgment

In this section, we extended our hypothesis from the control detection task, proposing that actively generated action plans are critical in shaping participants’ judgments. Similarly, we posited that the dynamics of these action plans serve as critical causal evidence informing their decisions. In the control judgment task, participants were tasked with moving a computer mouse freely while a single target stimulus was displayed on the screen. This resulted in a volitional and a displayed motor sequence in each trial.

In our approach, the encoder was employed to map volitional and displayed motor sequences to action plans, calculating both the Euclidean distances between these plans and their dynamics. This data served as predictor variables in a weighted logistic regression analysis, with leave-one-out cross-validation, allowing us to predict each trial’s judgment while accounting for individual differences in decision boundaries (Figure 5, more details see 5.7). Our trial-by-trial predictions tested whether the action plans and their dynamics could reliably predict judgments across various conditions and participants.

**Figure 5:**
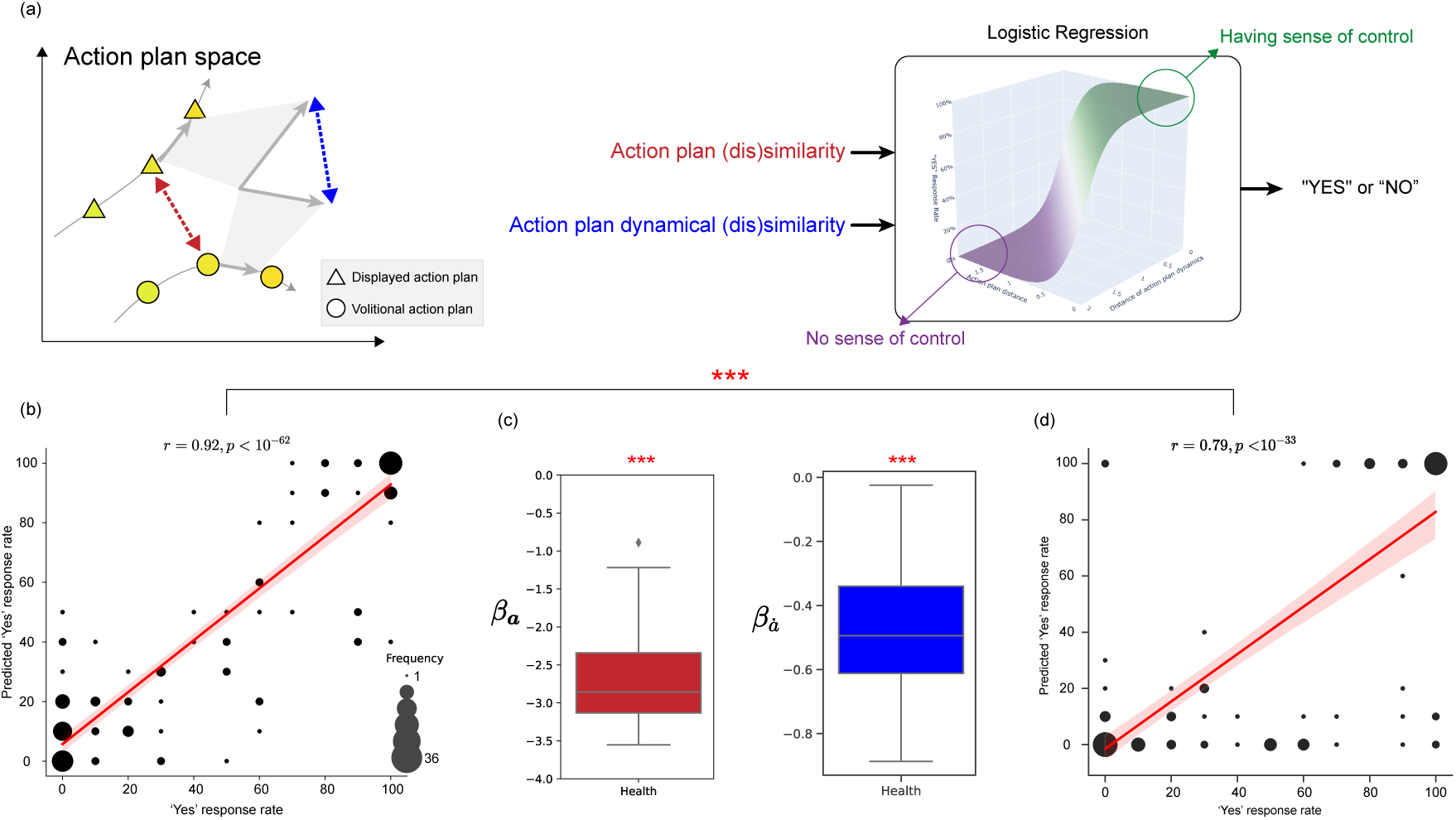
Predicted versus actual response rate in the control judgment task for the health group (a) Illustration of action plan space, highlighting how geometric distances and the dynamics of action plans are analysed through logistic regression to predict participants’ responses in the control judgment task. (b) The correlation between predicted and actual ’Yes’ response rates cross conditions and participants in the control judgment task indicating a strong alignment between the predictions and participants’ subjective judgments. (c) Logistic regression coefficients for action plans in the control judgment task. Presented are box plots contrasting the coefficients for the action plan distance (*β_α_*, left) and distance of action plan dynamics (*β_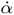_*, right). The coefficients are highly significantly below zero (*** *p <* 10*^−^*^10^), indicating that as the discrepancy between volitional and displayed action plans increases are less likely to feel in control during the control judgment task. (d) The predictive accuracy derived from micro-movements in control judgment tasks, while present, is significantly weaker compared to the robust correlation seen with action plans.

The results indicate a strong correlation (Figure 5b) between the predicted response rates and the actual response rates across conditions and participants (*r* = 0.92*, p <* 10*^−^*^62^). The high correlation suggests that the participants’ decisions were largely based on the geometric distance between the volitional and displayed action plans and the action plan dynamics.

The group-level prediction closely mirrors the actual response rates both qualitatively and quantitatively as illustrated in Figure S2a. These results suggest that the action plan contributes not only to control detection but also is involved in the judgment of self-relevant signals. This aligns with the initial qualitative observations showing consistency in actively using the similar action plan across two tasks within individuals.

#### 2.3.4 Evaluate the role of action plan dynamics in the control judgment task

To further explore the impact of the action plan and action plan dynamics on the response to the control judgment task, the fitted coefficients of the action plan and action plan dynamics from the logistic regression analysis were tested against 0 across participants. Both the coefficients of the action plan and action plan dynamics were found to be highly significant below 0 (*t*(24) = *−*19.63*, p <* 10*^−^*^15^ and *t*(24) = *−*11.79*, p <* 10*^−^*^10^, respectively, Figure 5c). The negative coefficients of the action plan and action plan dynamics indicate that the closer the geometric distance between the volitional and displayed action plans, as well as the action plan dynamics, the more likely participants were to recognise external stimuli as being caused by their actions. This highlights the significance of considering causal evidence in the sense of agency, as the action plan dynamics played a significant role in participants’ responses.

In conclusion, the results again highlight the significant influence of actively generated action plans and their intervention outcomes on the judgment of self-control over external stimuli. This emphasizes the critical role of active causal inference in shaping our sense of agency, suggesting that understanding the sense of agency requires a focus on active engagement and manipulation, i.e., intervention to the environment.

#### 2.3.5 The Predictive Power of Macroscopic Action Plans Over Microscopic Movements for Agency

In this study, we can consider action plans as macro-actions with respect to their spatial and temporal extent while instantaneous velocity sampled at a rate of 60Hz as micro-movement. Given that participants actively created action plans to infer their control capacity and form the sense of agency, and as evidenced by the preceding results, action plans served a critical functional purpose. We expected that their sense of agency would be more influenced by macroscopic evidence rather than microscopic motor-sensory contingencies. In other words, it’s crucial to investigate whether the sense of agency primarily emerges at a higher-functional level or simply stems from sensory-motor contingency.

To test this hypothesis, we analysed the effect of micro-movements on the sense of agency and compared it with the effect of action plans. Specifically, we conducted a similar analysis to that of action plans on micro-movements, using a cosine similarity distance metric that captures differences in the direction of micro-movement (see 5.8). This distance metric was chosen because the speed of volitional and displayed micro-movements remained constant due to the experimental design [for details see Oi et al., 2023]. By adopting this approach, we can gain insights into the contribution of micro-movements to the sense of agency.

##### The control detection task

The results (Figure 4f) revealed that micro-movements were not as effective as action plans in predicting accuracy in the control detection task in the healthy group (*r* = 0.47*, p <* 10*^−^*^9^). Although, the correlation between the predicted accuracy based on micro-movement and the actual accuracy was significant, the correlation coefficient was significantly lower than that of action plans (*z* = 8.33*, p <* 10*^−^*^15^). This finding suggests that action plans play a more prominent role in shaping the sense of agency than micro-movements in the control detection task.

##### The control judgment tas

In the control judgment task, the correlation (Figure 5d) between the predicted response rate based on micro-movement and the actual one was significant in the healthy group (*r* = 0.79*, p <* 10*^−^*^33^). However, comparing with the action plans, the correlation coefficients were significantly lower in both healthy (*z* = 4.55*, p <* 10*^−^*^5^). The results though micro-movement has significant predictive power in the control judgment task, it is not as effective as action plans in predicting the sense of agency.

#### 2.3.6 Active causal inference of Control Capacity through Action Plan Diversity

The results above have indicated that the action plan is a crucial element in the sense of agency. However, it remains uncertain whether participants would effectively use the action plan to infer their control capacity. Moreover, our results have revealed that the dynamics of action plans play a vital role in shaping the sense of agency, indicating that changing action plans may be an effective strategy for gathering more causal evidence. We hypothesised that participants can effectively use the action plan by actively increasing action plan diversity (Figure 6a,b). Specifically, we predicted that in low-noise environments, evidence accumulation would be rapid, leading to greater action plan diversity as participants change their action plans more frequently. Conversely, in high-noise environments, evidence accumulation would be slower, resulting in lower action plan diversity as participants change their action plans less frequently.

**Figure 6:**
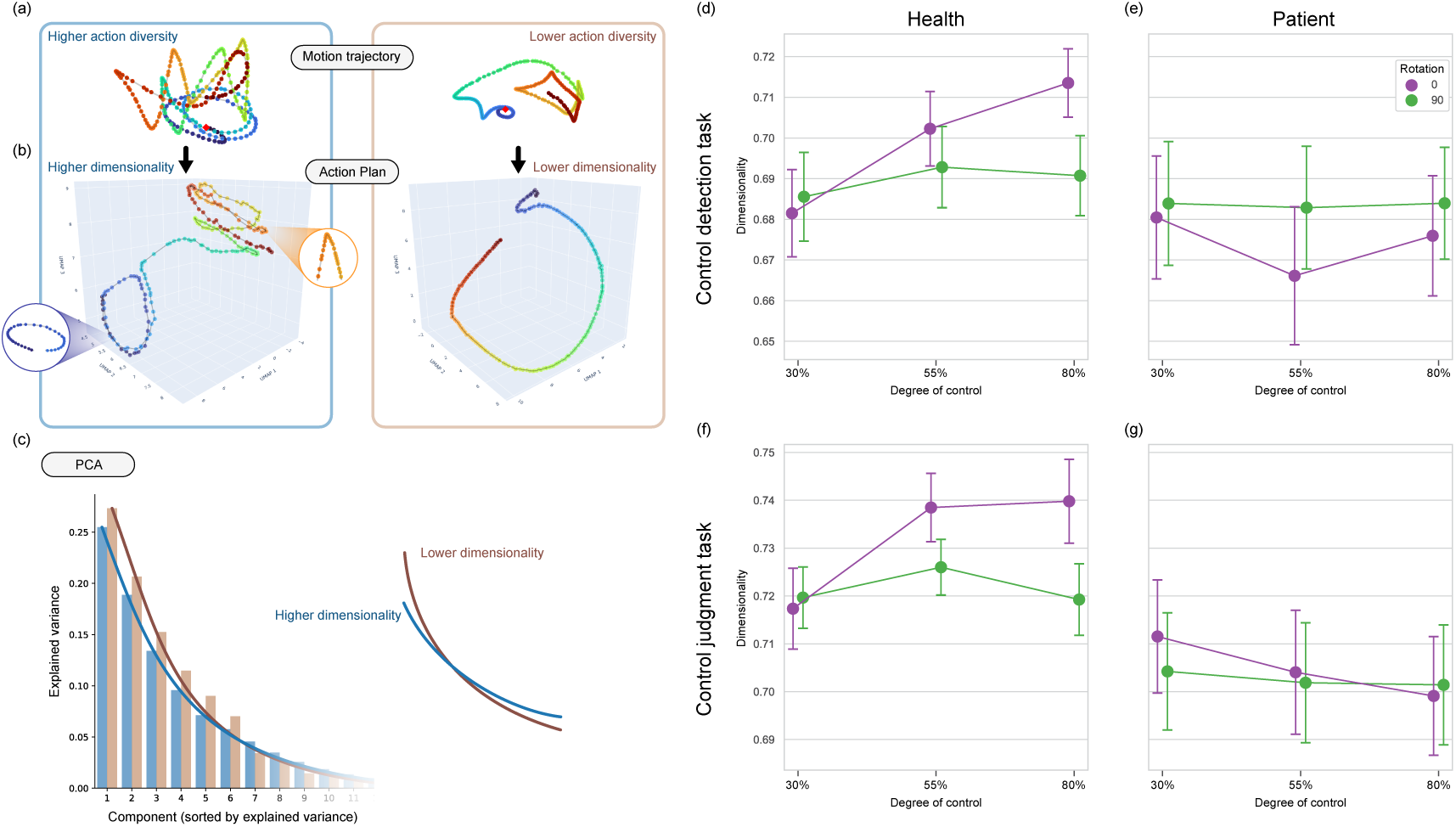
Figure delineates the trial-based diversity in action plans, essential for evidencing active causal inference. (a) Action plans exhibit trial-to-trial diversity. (b)The recruitment of diverse action plans necessitates involvement with various dimensions in the action plan space. This leads to differences in the dimensionality of the action plan distribution across trials. (c) In each trial, principal component analysis (PCA) was used on the action plan data to determine the distribution of explained variance by the principal components. When dimensionality is low, the explained variance is concentrated on a few principal components; conversely, high dimensionality leads to a more uniform distribution of explained variance. This variation in distribution can be captured by fitting it to an exponential distribution function. (d)(e) Action plan diversity measured by dimensionality across different degrees of control and rotation conditions in both healthy (d) and patient (e) groups in the control detection task. For the healthy group, a notable increase in action plan dimensionality with lower noise levels suggests that participants are actively gathering causal evidence within the action plan space, reflecting a robust sense of agency. The graph highlights the active exploration and manipulation of action plans, serving as a testament to the participants’ ability to explore and control actions within the space. Conversely, the patient group displays a relatively flat dimensionality profile, suggesting less active causal inference and a potential deficit in the sense of agency. (f)(g) This pattern was also observed in the control judgment task. Error bars denote standard error of the mean.

##### Measuring Action Plan Diversity through the Dimensionality

Given that changing the action plan necessitates the current action plan moving to a different position in the action plan space, which might require engaging different dimensions in the action plan space, we can consider that if the samples of action plan move and fall on a low-dimensional manifold, the action plan diversity is low. Conversely, if ones move and fall on a high-dimensional manifold, the action plan’s diversity is high.

To assess action plan diversity, we conducted principal component analysis on the action plan data in each trial and computed the explained variance distribution captured by the principal components (Figure 6c). Low dimensionality results in explained variance concentrated in a few principal components, while high dimensionality results in a more evenly distributed explained variance. We fitted the explained variance distribution to the function of exponential distribution and used the single fitted parameter *λ* to quantify dimensionality (see 5.9 for details).

##### Action plan diversity in the control detection task

The analysis of the action plan’s dimensionality in the control detection task employed a two-way ANOVA, focusing on the degree of control and rotation. In the healthy group, this analysis revealed a significant two-way interaction between the degree of control and rotation, as indicated by *F* (2, 48) = 6.48, *p* < .003. Furthermore, the main effects of both the degree of control and rotation were also significant, with *F* (2, 48) = 6.97, *p* < .002 and *F* (1, 24) = 5.93*, p* = .023, respectively. These results lend robust support to our hypothesis that the diversity of the action plan is greater under conditions of low noise and reduced in high-noise scenarios. This finding provides compelling evidence for the presence of active causal inference in the healthy group, as illustrated in Figure 6c.

##### Action plan diversity in the control judgment task

In a parallel investigation within the control judgment task, the impact of noise on action plan diversity was also assessed for each group. Here, the healthy group showed a significant interaction, marked by *F* (2, 48) = 4.22*, p* = .020, and notable main effects for both degree of control and rotation, with *F* (2, 48) = 3.90*, p* = .027 and *F* (1, 24) = 4.52*, p* = .044 (Figure 6f). These results align with the finding in the control detection task, reinforcing our understanding of active causal inference’s role in shaping the sense of agency under varying noise conditions.

In summary, for the healthy group, we observe a dynamic process where participants actively create and utilise action plans to assess their control capacity over the environment. By altering these action plans, they effectively gather causal evidence, aiding in the inference of their control capacity. Furthermore, the participants actively explore the action plan space, demonstrating an adaptive strategy to infer their control capacity across different noise levels in the environment.

### 2.4 Results from the Patient with schizophrenia group

In the patient group, the study explores how individuals with schizophrenia perceive, interact with, and adapt to their environment in terms of controllability and agency. This examination provides a distinct comparison to the healthy group in terms of qualitative and quantitative perspectives, highlighting how psychiatric conditions may influence the formation and utilization of action plans.

#### 2.4.1 Decreased Action Plan Diversity in Patient Group

The first noticeable distinction in the patient group, compared to the healthy group, is their notably lower action plan diversity in both control detection and control judgment tasks. This is evidenced by the lower dimensionality of their action plans, lacking the consistency observed in the healthy group across various conditions. The three-way ANOVA results confirm this result in the control detection task, showing a significant interaction between group, degree of control, and rotation (*F* (2, 96) = 5.15*, p* = .008), as depicted in Figure 6e. The simple main effect showed that, for the patient group, only the rotation’s main effect was significant (*F* (1, 24) = 6.94*, p* = .015). This pattern, especially when the rotation is 0 degrees and degree of control is 55%. However, the lack of a clear trend or tendency in this pattern suggests the possibility of a type I error, as the action plan diversity remains uniformly low regardless of the condition.

Similarly, in the control judgment task, a significant three-way interaction was found (*F* (2, 96) = 4.89*, p < .*01) (Figure 6g), but no significant interactions or main effects were observed for degree of control and rotation in the patient group. These findings suggest that the patient group’s action plan diversity is not altered by noise level, unlike the healthy group, and imply a potential difficulty in actively explorer action plans to harvest causal evidence.

#### 2.4.2 Exploring the Sense of Control through Action Plans in Schizophrenia

To further elucidate the manner in which individuals with schizophrenia utilise action plans in inferring controllability over their environment, we extended the same analytical approach to the patient group.

#### 2.4.3 The control detection task

As mentioned above, our observations in the control detection task highlighted a marked reduction in the diversity of action plans among this group. This diminished variety led us to anticipate a correspondingly lower detection accuracy compared to the healthy group, suggesting a potential impediment in the process of active causal inference within the patient cohort.

Consistent with these assumptions, the predicted detection accuracy for the patient group was indeed lower (Figure S3a), aligning with findings from one previous study [Oi et al., 2023], where patients with schizophrenia demonstrated notably poorer performance in control detection tasks than their healthy counterparts. However, it’s notable that the correlation between predicted accuracy and actual performance, though still significant (*r* = 0.515*, p <* 10*^−^*^10^), was much weaker compared to the healthy group.

Interestingly, this discrepancy predominantly stemmed from an underestimation of the detection accuracy in the patient group (Figure S3b). This underestimation suggests that the action plan analysis does not fully encapsulate the factors contributing to the patient group’s performance. It indicates the presence of additional compensatory strategies or elements used by these individuals, which aid them in achieving higher accuracy than what their action plans alone would predict.

##### Evaluate the role of action plan dynamics in the schizophrenic patient group

Following the result, we examined the role of action plan dynamics in the control detection task in the patient group. We found that compared with the healthy group, the correlation between predicted accuracy—derived without considering action plan dynamics—and the ground truth accuracy showed no significant drop (*z* = 1.25*, p* = 0.21). This result suggests that the causal and intervention evidence provided by action plan dynamics may not be utilised as effectively by the patient group as by the healthy group.

#### 2.4.4 The control judgment task

Despite the previously observed low diversity in action plans and diminished detection accuracy in this patient group, their response rates in the control judgment task were remarkably well predicted by the geometrical relationships of the action plan and its dynamics. This was evidenced by the high correlation between predicted and actual response rates (*r* = 0.90*, p <* 10*^−^*^55^), as shown in Figure S4a. The high correlation suggests that the participants’ decisions were largely based on the geometric distance between the volitional and displayed action plans and the action plan dynamics. This result also implies that, despite a lower sensibility in detecting sensory signals caused by themselves, individuals with schizophrenia can still adapt and optimise their decision criteria given the abnormally decreased action plans. This ability enables them to achieve a response profile level comparable to healthy individuals when assessing the level of self-control of a target using the action plan space. The finding is consistent with the previous study [Oi et al., 2023], where we found no significant difference between the two groups in the control judgment task.

Further analysis of their regression coefficients for the action plan and its dynamics showed no significant differences from the healthy group (*t*(48) = 0.61*, p* = 0.55 for the action plan and *t*(48) = 0.62*, p* = 0.54 for the dynamics, Figure S5). Yet, these coefficients were notably below zero (*t*(24) = *−*23.42*, p <* 10*^−^*^17^ for the action plan and *t*(24) = *−*17.42*, p <* 10*^−^*^14^ for the dynamics).

## 3 Discussion

### 3.1 Summary

This study unveiled a novel perspective on the sense of agency, uncovering its active component that had been previously overlooked. Our research shifted the focus from the conventional view of the sense of agency as merely a passive outcome of perceptual inference. We demonstrated that participants actively engaged in pattern-creation behaviour, employing stable, consistent, and idiosyncratic macro-action plans to counteract environmental noise for more accurate inference. Through generating causal evidence by intervention and monitoring the dynamics between their action plans and environmental changes, participants were able to more precisely infer their control capabilities, contributing significantly to their sense of agency. Most notably, our findings revealed that participants proactively expanded the diversity of action plans, thereby enhancing their exploration and accumulation of causal evidence, which in turn provided them with rich information for inferring control capacities and forming a sense of agency. Overall, this study provided compelling insights into the proactive components of the sense of agency, highlighting the importance of taking a more comprehensive and dynamic approach to understanding the conscious experience of agency.

### 3.2 Transformer-LSTM-based autoencoder

The transformer-LSTM-based autoencoder played a pivotal methodological role in this study, affording us the ability to effectively quantify action plans across larger spatial and temporal extents. Its success not only illuminated its ability of capturing abstract high-level motor commands but also showcased its potential allowing flexible experimental designs and data analysis for future research on the sense of agency. By affording greater flexibility in experimental design, we could shift away from overly restrictive task settings where the degree of freedom of action heavily influences the underlying sense of agency [Barlas and Obhi, 2013, Villa et al., 2020]. Moreover, without utilizing the deep artificial neural network, it would be difficult to deal with the highly non-linear nature of the motor control hierarchy [Winter, 2009, Chen et al., 2010]. Overall, this approach empowered us to design tasks with greater degrees of freedom, which is vital for understanding the active nature of the sense of agency.

### 3.3 Active causal inference and the sense of agency

The current study posits that the sense of agency as an inference outcome of active causal inference process. Although this study does not strictly adhere to the mathematical formulation of active inference within the framework of the free energy principle, we believe our findings resonate with the core notions of active inference [Friston, 2010a]. The causal relationship between our intentions and subsequent changes in sensory feedback and motor output are obscured by uncertainty. Consequently, this study provides an interactive and dynamic perspective on the sense of agency. By forming action plans and using internal generative models to predict the outcomes of executing these plans, individuals engage in a form of hypothesis testing [Seth, 2016, Donnarumma et al., 2017, Friston et al., 2012]. This interactive process is crucial in avoiding the illusionary sense of control that may arise from solely correlational evidence.

In this way, the study provides a more holistic view of SoA, potentially integrating and reconciling various existing perspectives. The inferential nature of SoA aligns with both retrospective perspective including the theory of apparent mental causation [Wegner and Wheatley, 1999, Wegner, 2003] and the Bayesian inference perspective [Moore and Fletcher, 2012, Yano et al., 2020, Wen and Imamizu, 2022, Legaspi and Toyoizumi, 2019]. The inference process necessitates a generative model for making predictions about outcomes and computing prediction errors to update current inferences. In this context, the comparator model, particularly the forward model [Frith et al., 2000, Blakemore et al., 1999], becomes an integral part of the inference process, functioning as the generative model. Finally, active inference plays a pivotal role in generating action policies aimed at maximizing information gain and minimizing uncertainty [Friston et al., 2015], thereby optimizing the inference process. This active engagement in shaping one’s sense of agency reflects a deeper understanding of how individuals interact with and perceive their control over their actions and the environment.

### 3.4 Exploration and exploitation

The active causal inference process in our study can be seen as a form of exploration in action space [Wen et al., 2020a]. When individuals increase the diversity of their action plans, they are effectively searching the actionable space. People draw from diverse action plans when exploring control, and the sense of agency is greatly affected by the diversity of action plans in such exploration. It is evident that the exploration of action space and understanding our own control capacity is crucial for our survival and development, as demonstrated in developmental studies on the reinforcement of sensory feedback on infant behaviours [Rochat and Striano, 1999, Rovee and Rovee, 1969]. Indeed, exploration in action space is connected to the internal reward system as one type of intrinsic motivation [Karsh and Eitam, 2015, Karsh et al., 2016].

The focus of our study was primarily on examining active inference behaviour in tasks specifically designed to assess participants’ ability to infer controllability over environmental stimuli. This objective aligns closely with exploratory behaviours, wherein participants engage in varied action plans to probe and understand their control capacity. However, in daily life, our objectives often encompass more than just exploratory aims. Many of our actions are directed towards specific, goal-oriented objectives, necessitating exploitation behaviours. The delicate balance between exploration and exploitation, as highlighted in previous research [Cohen et al., 2007, Schwartenbeck et al., 2013, Kaplan and Friston, 2018], is a critical aspect of behavioural regulation to adapt their living environment. To enhance the ecological validity of our findings, future research should explore how the objective of control capacity inference intertwines with other task objectives [Schwartenbeck et al., 2019]. Such research should consider how different tasks impose constraints on the action space, potentially limiting action diversity and influencing the process of inferring control capacity. This broader approach would offer a more comprehensive view of how we navigate and control our environment.

#### 3.4.1 Implications for the schizophrenia patient group

The findings of the current study shed new light on the sense of agency in schizophrenia patients. The research revealed that individuals with schizophrenia demonstrated decreased action plan diversity in all experimental conditions, indicating limited and rigid capacity to infer control ability actively. This marked difference between the healthy and patient groups highlights the challenges faced by individuals with schizophrenia. However, despite their lower action plan diversity, the patient group still displayed a similar behavioural pattern to the healthy group in the control judgment task. It is plausible that the patient group has adapted their self-other decision criteria (or exaggerated “prior” in the Bayesian sense) based on their physical condition, providing a reasonable explanation for this outcome. Despite this, the action plan analysis was still effective in predicting the patient group’s responses in the control judgment task, indicating that they follow the same principle of using action plan space for inference (i.e., the geometric relationship).

It is important to note that control judgment is subjective and can be calibrated by adjusting the prior. In contrast, control detection requires the capacity to identify self-relevant signals from the noise. The predicted accuracy based on the patient group’s action plan data was slightly lower than the actual accuracy. This discrepancy could be due to the fact that the patient group may have developed other ways to detect self-relevant cues, not considered in the current study, to compensate for their reduced action plan information for control detection.

The difference between the healthy and patient groups in the control detection task and action plan diversity can potentially be used to develop new diagnostic markers for schizophrenia. Furthermore, our new approach can also be used to study the sense of agency in patients with schizophrenia in a more comprehensive way. Overall, these findings highlight the importance of examining the sense of agency in schizophrenia patients and suggest that action plan analysis can provide a valuable methodology for understanding abnormality in their sense of agency.

## 4 Conclusion

This study reveals a dynamic aspect of the emergence of sense of agency (SoA), highlighting its active component beyond passive perceptual inference. Participants actively formed macro-action plans to counteract environmental noise, a process effectively captured using a transformer-LSTM-based autoencoder. This approach allowed for the accurate prediction of task performances in control detection and judgment tasks. Notably, schizophrenia patients showed reduced action plan diversity, suggesting impaired active causal inference in both qualitative and quantitative perspectives.

Overall, our results highlight the importance of active engagement in shaping SoA, suggesting a dynamic interaction with the environment. This contributes to a deeper understanding of SoA in both healthy individuals and those with psychiatric conditions. Moreover, specific manifestations in schizophrenia would support to diagnose and investigate pathophysiology of the illness.

## 5 Methods

The dataset employed in this research was sourced from one preceding study, as detailed in [Oi et al., 2023]. Comprehensive descriptions of the experimental protocols, design, and statistical data on the patient population involved.

### 5.1 Participants

This study included 26 individuals (nine females and sixteen males, aged 26–56 years) diagnosed with schizophrenia and 27 healthy individuals (nine females and sixteen males aged 24–58 years) as participants. Patients with schizophrenia were recruited from Keio University Hospital and Komagino Hospital in Tokyo, Japan. The study was approved by the ethics committees of Keio University Hospital, Komagino Hospital, and the University of Tokyo. One patient was also excluded due to withdrawing their consent. Additionally, two healthy participants were excluded because they attributed nearly all of their responses to self in the control judgment task, suggesting they did not understand the task. This resulted in a final sample of 25 patients and 25 age, sex, and dominant hand-matched healthy controls.

Patients with schizophrenia who met the diagnostic criteria outlined in the Diagnostic and Statistical Manual of Mental Disorders-5 (DSM-5). The severity of their symptoms was evaluated using the Positive and Negative Syndrome Scale (PANSS) (more details see Oi et al. [2023]).

### 5.2 Behavioural Tasks

#### 5.2.1 The general task design

In this study, we designed tasks in which participants used a computer mouse to control the movement of a *target dot* for a duration of 5 seconds. Two types of interference on the moving direction of the target were introduced (see Figure 1cabala): degree of control and affine transformation (rotation). The degree of control (30%, 55%, or 80%) was varied by altering the proportion of the participant’s mouse movement that influenced the dot’s moving direction. For example, in the 80% control condition, the dot’s moving angle was determined by 80% of the participant’s mouse movement and 20% of a predetermined movement trajectory. The affine transformation was manipulated by adding a 0- or 90-degree rotation to the direction of the dot’s movement [Wen and Haggard, 2020, Wen et al., 2020c, 2018]. This resulted in six different conditions, each of which was repeated 10 times for a total of 60 trials per task.

#### 5.2.2 The control detection task

In the control detection task (Figure 1a), participants were presented with a screen displaying three dots, one of which was designated as the target dot. The movement of the target dot was partially under the control of the participant via their mouse movements, while the other two dots followed pre-determined paths. After a brief 5-second period, the three dots remained on the screen, and participants were asked to identify and select the target dot.

#### 5.2.3 The control judgment task

In the control judgment task (Figure 1b), participants were asked to decide whether they felt they could control the moving direction of a dot (target) on the screen after freely moving the mouse for 5 seconds. The dot began at the centre of the screen and started moving once the participant began moving the mouse. The algorithm used to generate the dot’s motion was the same as the one used for the target dot in the control detection task. Participants responded by saying “yes” or “no” in Japanese.

### 5.3 Data Recording

#### 5.3.1 Motion trajectory and motor sequence

In every trial, participants moved a computer mouse for five seconds at a sampling frequency of 60 Hz. This resulted a two-dimensional (horizontal and vertical) trajectory of mouse cursor movement, called *motion trajectory* ***P*** = (***p***_1_, ***p***_2_*, · · ·,* ***p****_n_*), where ***p****_t_* = (*p_x,t_, p_y,t_*) *∈* ℝ^2^ represents the cursor’s position at the *t*-th time point. A *motor sequence*, ***M*** = (***m***_1_, ***m***_2_*, · · ·,* ***m****_n−_*_1_), where ***m****_t_* = (*m_x,t_, m_y,t_*) *∈* ℝ^2^ is defined as the temporal difference between two consecutive time points of the motion trajectory, where **m_t_** = ***p****_t_*_+1_ *−* ***p****_t_* represents the velocity vector of the cursor movement at time-step *t*. The recorded motor sequence values (in pixels) were normalised by dividing them by the width of the presentation screen resolution (in pixels) in all the following analyses.

#### 5.3.2 Volitional and displayed sequences

In this article, we use the term *volitional* to refer to the data obtained from participants’ unaltered movements, which is denoted with the left-subscript *_vol_*(.), such as *_vol_****M*** and *_vol_****P*** . On the other hand, we use the term *displayed* to refer to the data that had undergone experimental interference and was subsequently presented on the screen. The data is denoted with the left-subscript *_disp_*(.), such as *_disp_****M*** and *_disp_****P*** .

### 5.4 Transformer-LSTM-based Autoencoder

To capture high-level, abstract, and low-dimensional motor commands (i.e., the action plans), we employed an autoencoder trained to learn abstract action plans from **volitional** motor sequences generated by each participant. The architecture of the autoencoder consisted of a transformer-based encoder, a hidden layer with 16 neurons, and a long short-term memory (LSTM) decoder (as depicted in Figure 7).

**Figure 7:**
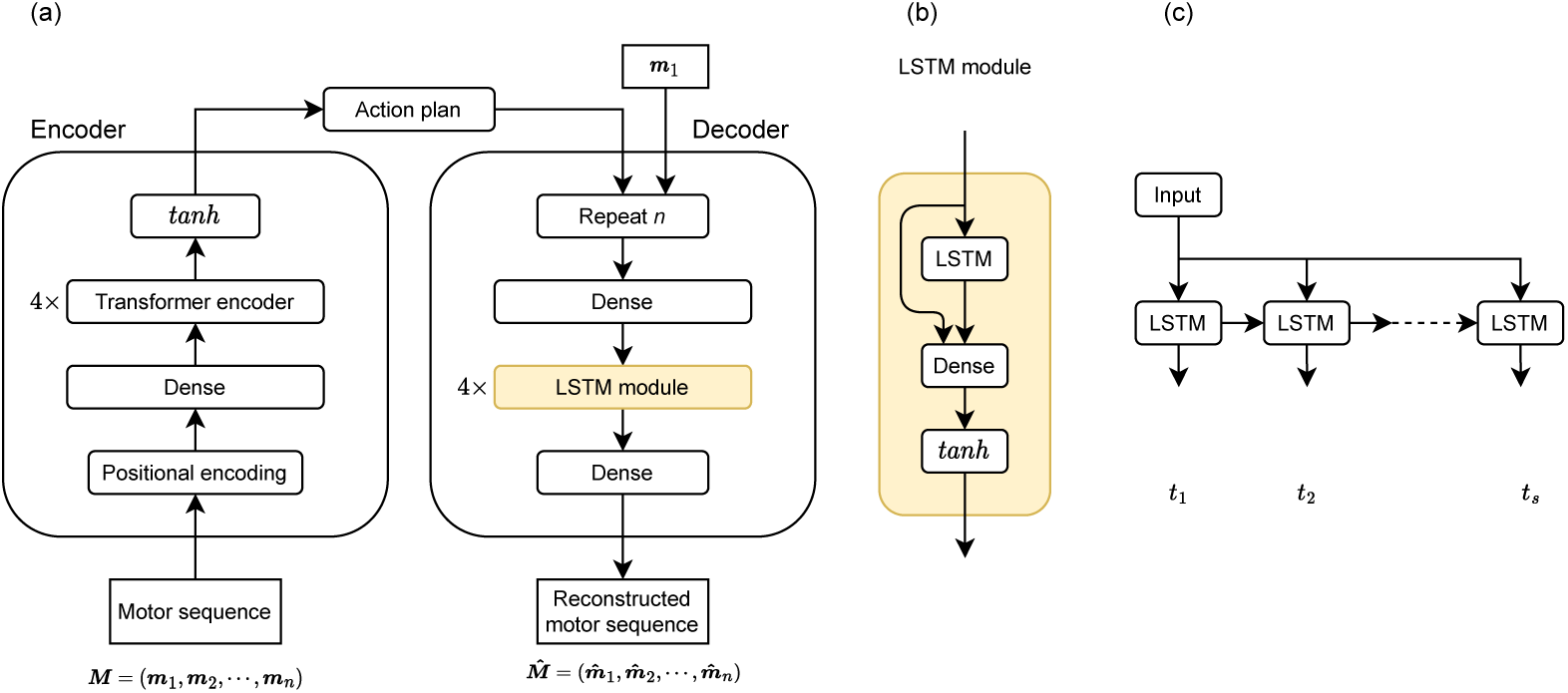
Transformer-LSTM-based Autoencoder. The transformer-LSTM-based autoencoder is composed of three main components: a transformer-based encoder, a bottleneck, and a LSTM-based decoder. The encoder takes a motor sequence as input and maps it to an action plan. The bottleneck, i.e., the action plan space, serves as a compression layer to capture the essential features, i.e., the action plans, of the input motor sequence. The decoder then takes the action plan and the first time point of the motor sequence as inputs and reconstructs the original motor sequence. (b) The structure of the LSTM module is illustrated in yellow in (a). (c) The temporal unfolding of the LSTM unit. At each time point, the LSTM unit receives the same higher-level motor command. Therefore, the higher-level motor command and the current internal state of the LSTM unit co-determine the output at each time point.

#### Encoder

The transformer encoder (encoder only architecture, see Vaswani et al. [2017]), represented by a function *f_enc_*(*·*), consisting of a stack of 4 transformer blocks, each comprising a self-attention layer with 32 heads and a feed-forward layer with 64 dimensions, is utilised to capture the long-range dependencies of motor sequences. To clip the output values remain within the range of [*−*1, 1], a hyperbolic tangent activation function was applied to the final output of the transformer encoder. Mathematically, the encoder can be represented as follows:

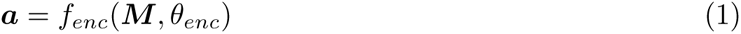

Where ***M*** denotes a motor sequence, ***a*** denotes the corresponding action plan, and *θ_enc_* denotes the learnable parameters of the encoder, which were optimised during training.

#### Positional Encoding

The position encoding is mimicking the positional encoding in the Transformer architecture Vaswani et al. [2017]. The position encoding is a matrix of sine and cosine values with varying frequencies. The position encoding is calculated as follows:

Let *n* be the length of the input motor sequence, *d* = 16 be the number of dimensions used to encode position information, and *l_max_* = 300 be the maximum possible length of the input sequence. We can then define the following matrix:

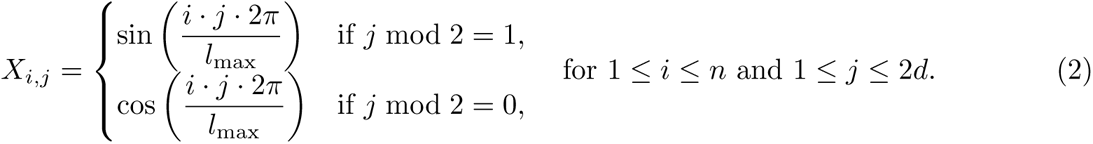

This gives us a row of the matrix represents a different time step, and the columns represent the sine and cosine values at different frequencies. The position encoding is concatenated to the feature dimension of the input of the Transformer encoder to capture the temporal information of the motor sequence.

#### Hidden layer (action plan space)

The encoder outputs a 16-dimensional hidden embedding, called an *action plan* ***a*** = (*a*_1_*, a*_2_*, · · ·, a*_16_) *∈* R^16^, which is used to represent high-level, abstract, and low dimensional motor commands generating motor sequences. The domain of the action plan is defined as *A* called *action plan space*.

#### Decoder

The decoder, represented by a function *f_dec_*(*·*), maps the action plans back to the original dimension. To mimic the generative process from the abstract motor command (action plan) to motor sequences, a stack of LSTM modules were utilised. The LSTM module is designed with a structure similar to the ResNet architecture [He et al., 2016] to mitigate the problem of vanishing gradients. The LSTM module is depicted in Figure 7b. The decoder stack 4 LSTM modules, each comprising a hidden layer with 64 neurons. The LSTM module is trained to reconstruct the motor sequence ***M*** given the action plan ***a*** and the first velocity vector ***m***_1_ of the motor sequence ***M*** . The supply of the first velocity vector is motivated by the focus on the action plan, rather than the initial condition of the motor sequence. The decoder outputs a reconstructed motor sequence ***M̂*** with the same dimension as the input motor sequence ***M*** . Mathematically, the decoder can be represented as follows:

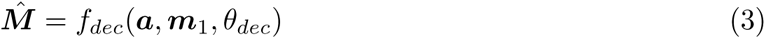

*θ_dec_* denotes the learnable parameters of the decoder, which are optimised during training.

#### 5.4.1 Loss function

The mean squared error (MSE) between the input motor sequence *M* and its reconstruction *M̂* is used as the loss function during training. The MSE is mathematically defined as:

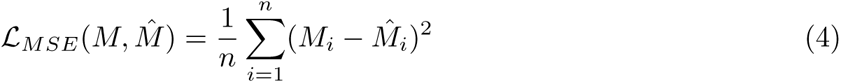

where *n* is the number of elements in the motor sequence.

#### 5.4.2 Training

The goal of the training process is to learn to embed motor sequences of 1 second length (60 time points) into a 16-dimensional action plan embedding. To achieve this, a 1-second (60 time point) rolling window is used to extract **volitional** motor sequences for each trial, resulting in 241 1-second motor sequences for each trial.

To further improve the training process, a secondary step is performed during training. For each 1-second motor sequence, a consecutive 20 to 60 time points (i.e., a 0.33 to 1 second subsequence) were randomly extracted to form motor sequences of variable length. Our preliminary experiments have shown that training with variable length motor sequences results in better generalisation and effectively decreases the validation loss compared to training with fixed length motor sequences.

The mini-batch size used during training is 64, with the learning rate starting from 10*^−^*^3^ and the weight decay set to 10*^−^*^4^. The Adam (adaptive moment estimation) [Kingma and Ba, 2014] optimiser is used for training.

#### 5.4.3 Validation

In one training epoch, once all training samples have been processed, the validation loss is calculated using a set of validation samples. During validation, the autoencoder uses 1-second motor sequences as input without the additional step of sub-sequence sampling.

#### 5.4.4 Early stopping criterion

Both the training and validation loss are calculated after each epoch. To prevent overfitting, an early stopping criterion is implemented. Training was stopped when the validation loss stops decreasing for 20 consecutive epochs, as this is a commonly used early stopping criterion to prevent overfitting. This criterion ensures that the model is not trained for too long, reducing the chances of overfitting and increasing the chances of generalisation.

During training, we discovered that two models had significantly higher validation losses. To address this issue, we retrained the models using different random seeds for the training-validation trial split. This resulted in the successful lowering of validation losses. All models completed training between 100 and 200 epochs. Models and training were implemented using the PyTorch library [Paszke et al., 2019].

### 5.5 Action plan analysis

#### 5.5.1 Mapping motor sequences to action plans

To conduct our analysis, it is essential to extract high-level action plans from recorded motor sequences. This involves slicing a motor sequence into smaller subsequences using a sliding window approach. Here, we used a window size *w* = 60 and step size *s* = 1. The slicing process results in *n − w* + 1 subsequences, where *n* is the length of the motor sequence. Next, we used an encoder to map each subsequence to an action plan, which resulted in a sequence of action plans denoted by ***A*** = (***a***_1_, ***a***_2_*, · · ·,* ***a****_n−w_*_+1_). In this sequence, ***a****_i_* refers to the action plan of the *i*-th motor subsequence.

#### 5.5.2 Action plan dynamics

We define *action plan dynamics* ***ȧ*** *_t_* as the difference between two consecutive temporal action plans ***a****_t_*_+1_ and ***a****_t_*. This difference is expressed mathematically as follows:

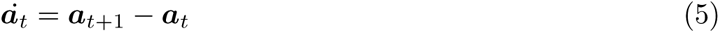

#### 5.5.3 Distance metric in action plan analysis

In this study, Euclidean distance was used as the distance metric to compare two action plans ***a*** and ***a****^′^*

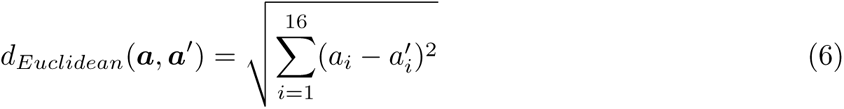

Here, *a_i_*and 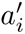 denote the *i*-th element of action plans ***a*** and ***a****^′^*, respectively.

### 5.6 Predicting the behaviour profile of the control detection task

In each trial, a one-second sliding window was applied to the volitional and the three displayed motor sequences to obtain a series of one-second subsequences (Figure 2). The action plan of each subsequence was computed by running the encoder on the subsequence. At each sample time point of the action plan sequence, we calculated the distance between the volitional action plan and the displayed action plans. This results in three distance series. We also applied the same computation to the action plan dynamics of the volitional and displayed action plans (Figure 3a).

#### Compute one-vs-rest AUCs

To predict the decision process of participants in each trial of the control detection task, we employed the one-vs-rest receiver operating characteristic (ROC) curves to quantify the separability of three distance series [Hanley and McNeil, 1982]. The areas under curve (AUC) were then calculated for each ROC curve to provide a scalar measure of the performance of each distance series. We repeated this computation procedure for both the action plan and action plan dynamics, resulting in six AUC values in total (Figure 3b). The distance series with the highest AUC among the six was selected as the most salient self-relevant feature, enabling us to predict the participants’ response in each trial. The computation of ROC curves and AUC was done using the scikit-learn library [Pedregosa et al., 2011].

#### Decision criterion

Let *AUC_a,_*_1_*, AUC_a,_*_2_*, AUC_a,_*_3_ denote the areas under curve for the receiver operating characteristic (ROC) curves corresponding to the distances between the volitional action plan and displayed action plans, and let *AUC_ȧ_*_,1_*, AUC_ȧ_*_,2_*, AUC_ȧ_*_,3_ denote the areas under curve for the ROC curves corresponding to the distances between the volitional action plan dynamics and displayed action plan dynamics. The index of the largest area under curve from either the action plan distances or action plan dynamics, *i^∗^*, is selected as the stimuli with the most salient self-relevant feature, i.e.,

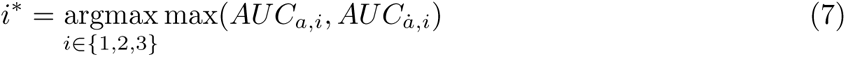

Therefore, the stimulus *i^∗^* is the one that is most likely to be selected as the target by the participant and was considered as the simulated decision that participants made in the control detection task.

### 5.7 Predicting responses in control judgment task

In the control judgment task, participants freely move a computer mouse and a single stimulus (target) was displayed. This results in one volitional and one displayed motor sequence in each trial. The two motor sequences were mapped to action plans using the encoder. Similar to the process in the control detection task, the distances between the volitional and displayed action plans so as the action plan dynamics were calculated. However, unlike the control detection task, the distances from all sample time points in a trial are averaged to obtain a single value for both the action plan and its dynamics

Participants were asked to make a binary decision (i.e., either “yes” or “no”) at the end of a trial in the control judgment task. By incorporating the information from the action plan and the action plan dynamics into the decisions of control judgment, the distances should be able to effectively predict the decisions of participants on a trial-by-trial basis.

#### Logistic Regression Analysis with Leave-One-Out Cross-Validation

If participants made their responses using action plans, we can predict their responses by analysing the geometrical relationships between the action plans. To test this hypothesis, we conducted a leave-one-out cross-validation of a weighted logistic regression analysis. We used the averaged distances of the action plan and its dynamics as two predictors, and the participant’s binary response (denoted as *ψ_i_*) as the dependent variable, where *ψ_i_* can have a value of 1 (indicating a “yes” response) or 0 (indicating a “no” response).

For each participant, we predicted their response in each trial by fitting a weighted logistic regression model to the remaining trials. The logistic regression model is defined as:

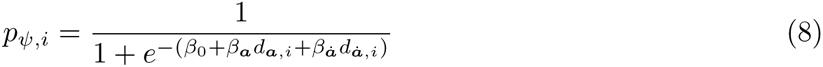

Where *p_ψ,i_*is the probability of making a response of 1 at trial *i*. *d****_a_****_,i_* and *d****_ȧ_****_,i_* denote the averaged distances between the volitional and displayed action plans and the action plan dynamics at trial *i*, respectively. *β*_0_, *β****_a_***, and *β****_ȧ_*** are model free parameters.

Finally, a decision function *f_ψ_* is defined as:

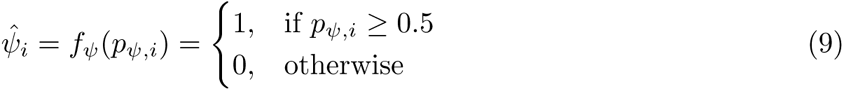

Therefore, the predicted response 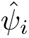 at trial *i* can be obtained by applying the decision function *f_ψ_* to the probability *p_ψ,i_*.

To address the class imbalance problem, the weight assigned to each trial *i* is proportional to the inverse of the frequency of its corresponding response in the training set. Specifically, let *ψ_i_* denote the response of in trial *i*. Then, the weight *w_i_* for trial *i* is defined as:

To address the class imbalance problem, we assign weights to each trial based on the frequency of its corresponding response in the training set. Let *N* be the total number of trials, *n*_1_ be the number of trials with responses of 1, and *n*_0_ be the number of trials with responses of 0. Then, for each trial *i*, the weight *w_i_*is defined as follows:

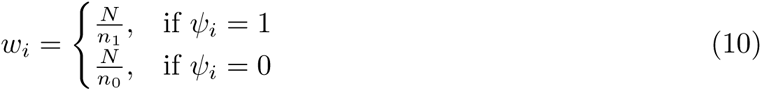

Thus, trials with less frequent responses are assigned greater weight to ensure their impact is not overshadowed by more common responses. Using these weights, the logistic regression objective function becomes:

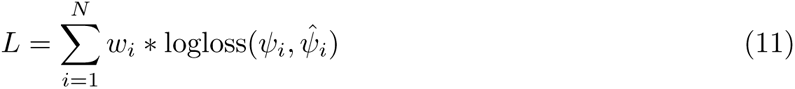

where logloss is the logistic loss function.

To test whether the coefficients are significantly different from 0 at group level, we use full trials to fit weighted logistic regression for each subject and obtained a single value for each coefficient. We next used a one-sample t-test to test whether the coefficient is significantly different from 0.

This logistic regression analysis was conducted using the scikit-learn library [Pedregosa et al., 2011].

### 5.8 Micro-movement analysis

To determine whether action plans at a macro-scale hold more explanatory power than transient micro-movements, we undertook a micro-movement analysis. This analysis defines micro-movement as a single velocity vector sample ***m****_t_*, taken at a 60Hz sampling rate. *t* denotes the time index of the velocity vector sample. Comparable to action plan analysis, micro-movement analysis also considers the similarity of velocity vectors and their dynamics, i.e., the acceleration ***ṁ*** = ***m****_t_*_+1_ *−****m****_t_* of movement.

#### Distance metric in micro-movement analysi

Our experimental design ensures that all moving objects have the same speed, with only differences in moving direction. Thus, cosine similarity is an appropriate metric to quantify the difference between two velocity vectors. Cosine distance based on cosine similarity between two velocity vectors ***m*** and ***m****^′^* is defined as:

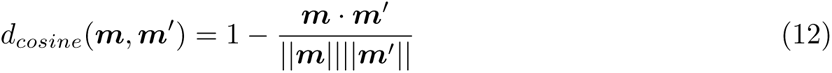

### 5.9 Measuring action plan diversity via dimensionality

In this study, we considered dimensionality as a proxy to assess the diversity of action plans. In the action plan space, a point represented an action plan. To make a different action plan implied activating different neurons in the action plan space, which was equivalent to recruiting more dimensions in the action plan space. Therefore, by measuring the dimensionality in the action plan space, we could quantify the diversity of action plans.

In each experimental trial, we performed Principal Component Analysis (PCA) on the action plan dataset ***A***, and subsequently computed the distribution of explained variance. We then fitted the distribution of explained variance to the probability density function of exponential distribution *f* (*x*; *λ*) defined for *x ∈* ℝ as:

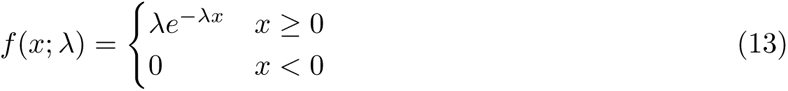

Where the rate parameter *λ* is determined through fitting. Finally, we define the dimensionality *dim* as:

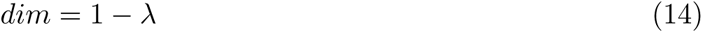

#### Analysis of Variance (ANOVA) on dimensionality

To test whether the dimensionality of action plans is significantly different across conditions and groups, we computed the average dimensionality of action plans across all trials for each condition and participant. We then conducted a three-way ANOVA on the averaged dimensionality of action plans with two within-subject factors: degree of control (30%, 55%, and 80%) and rotation (0 and 90 degrees) and one between-subject factor: group (healthy and patient). Once the three-way interaction was significant, we performed a two-way ANOVA with within-subject factors: degree of control and rotation for each group.

## Author contributions

A.C. and W.W. conceptualized the research goals and framework, with A.C. developing the study’s methodology and analysis approach. A.C. generated the software and undertook the formal analysis of the data. Data collection and management were overseen by H.O. and T.M. The initial manuscript draft was penned by A.C. with critical review and editing by all authors, A.C., H.O., T.M., and W.W. Project administration and funding acquisition were orchestrated by W.W. and T.M. All authors have given their approval for the final manuscript to be published.

## Acknowledgments

This study was supported by the JST FOREST Program (Grant Number JPMJFR2144) and the Japan Society for the Promotion of Science (KAKENHI 19H05729, 21H03780, and 22H04781).

## Appendix A Glossary of Notations

ℝ: Real number
ℕ: Natural number
M: Motor sequence
M̂: Reconstructed motor sequence
M: Motor sequence space
m: Velocity vector
P: Motion trajectory
p: *_t_ -* Mouse position at time *t*
a: Action plan
A: Action plan sequence
ȧ: Action plan dynamics
Ȧ: Action plan dynamics sequence
A: Action plan space
a_i_: The activation of the *i*th neuron of the action plan ***a***

## 6 Supplementary Materials

**Figure S1:**
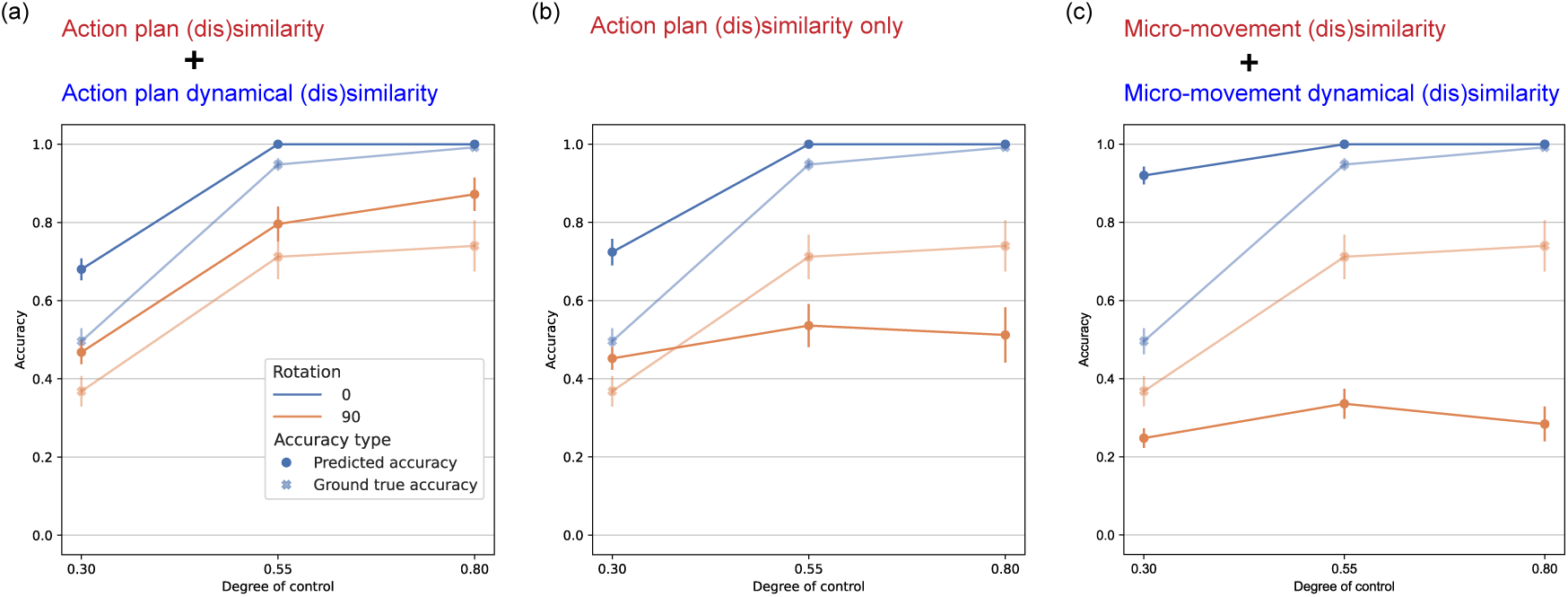
Group-level predicted versus actual task performance in the control detection task for the health group (a) Group-level analysis showing how predicted accuracies align with actual accuracies, reflecting the influence of two types of noise on control inference. (b) The absence of action plan dynamics in predictions leads to a qualitative mismatch with the actual group behavioural profile, reinforcing the premise that action plan dynamics provide essential causal evidence for precise control inference. (C) Demonstrates a group-level analysis showing the inability of micro-movement predictions to accurately capture the trend in actual accuracies across control conditions. Error bars denote standard error of the mean.

**Figure S2:**
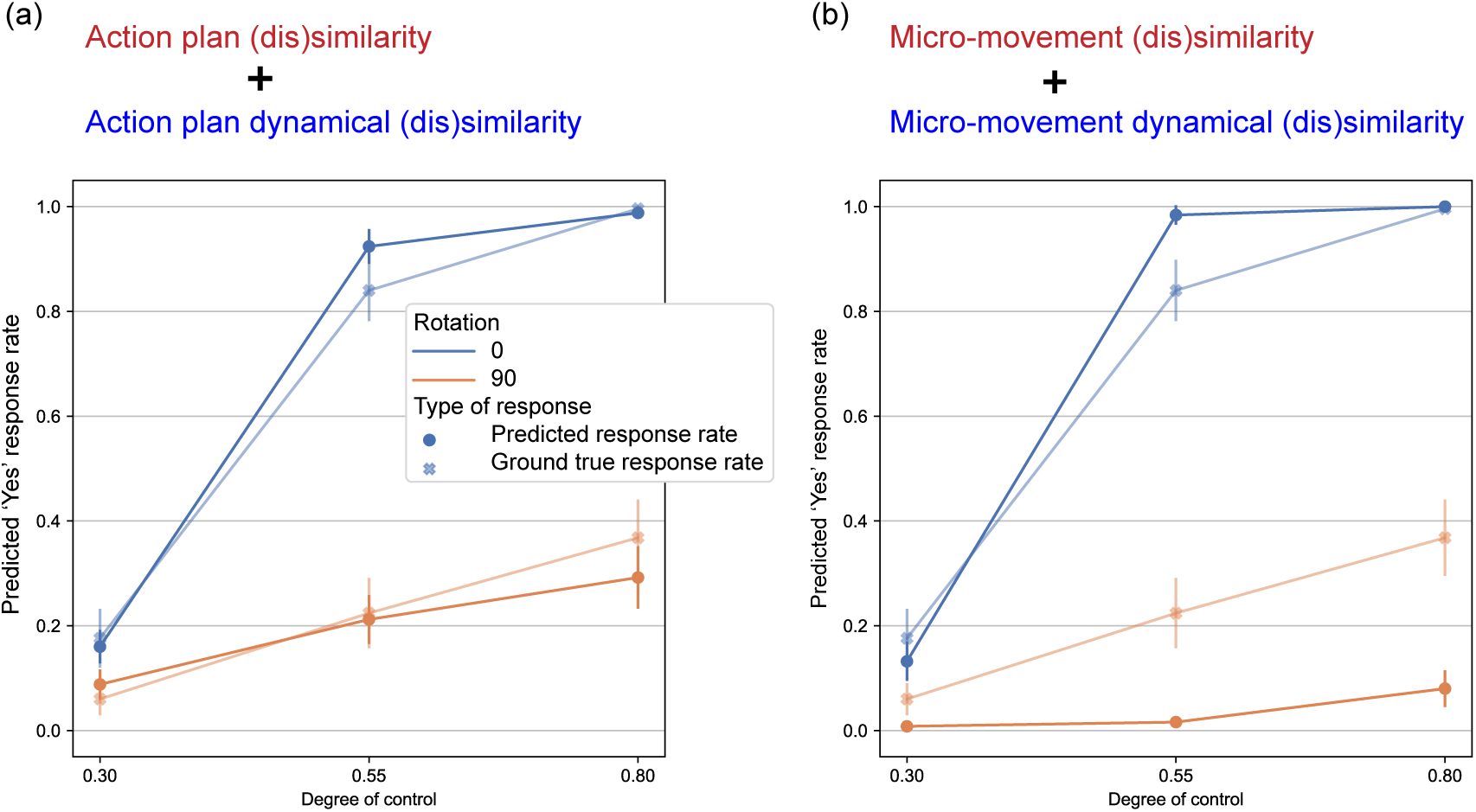
Group-level predicted versus actual task performance in the control judgment task for the health group (a) The group level response aggregating the predictive accuracy and the ground truth accuracies in each condition. Error bars denote standard error of the mean.(B) Reveals at the group level, where predicted response rates fail to qualitatively align with actual rates, suggesting a gap in the micro-movement predictive approach in reflecting group-wide behavioural patterns.

**Figure S3:**
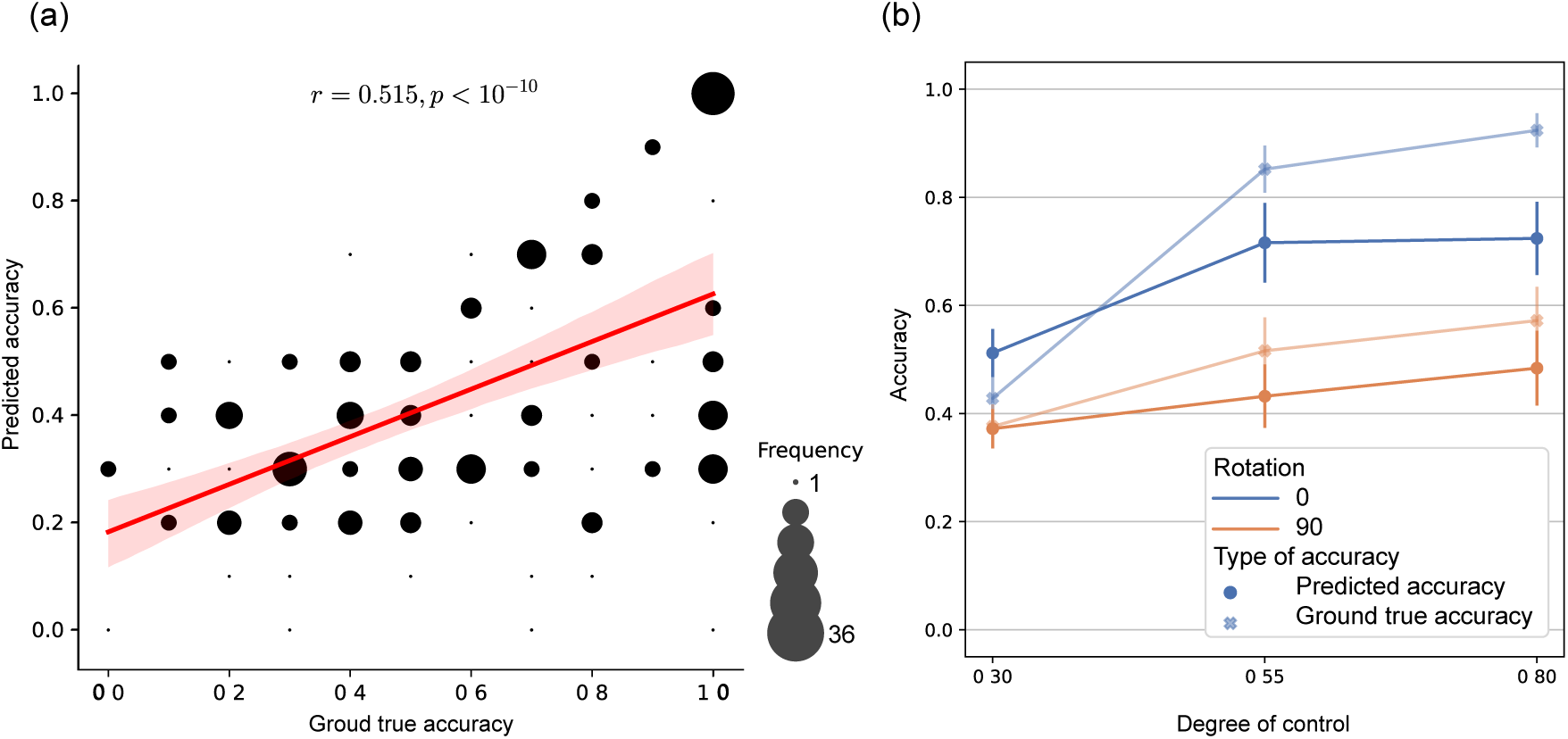
Correlation between predicted and ground truth accuracy in the control detection task in the schizophrenic patient group. (a) Illustrates the correlation between predicted and ground truth accuracy, with point size indicating the frequency of data points. (b) Presents a comparative analysis of predicted and actual accuracies across conditions, with a marked underestimation of detection accuracy visible in the patient group.

**Figure S4:**
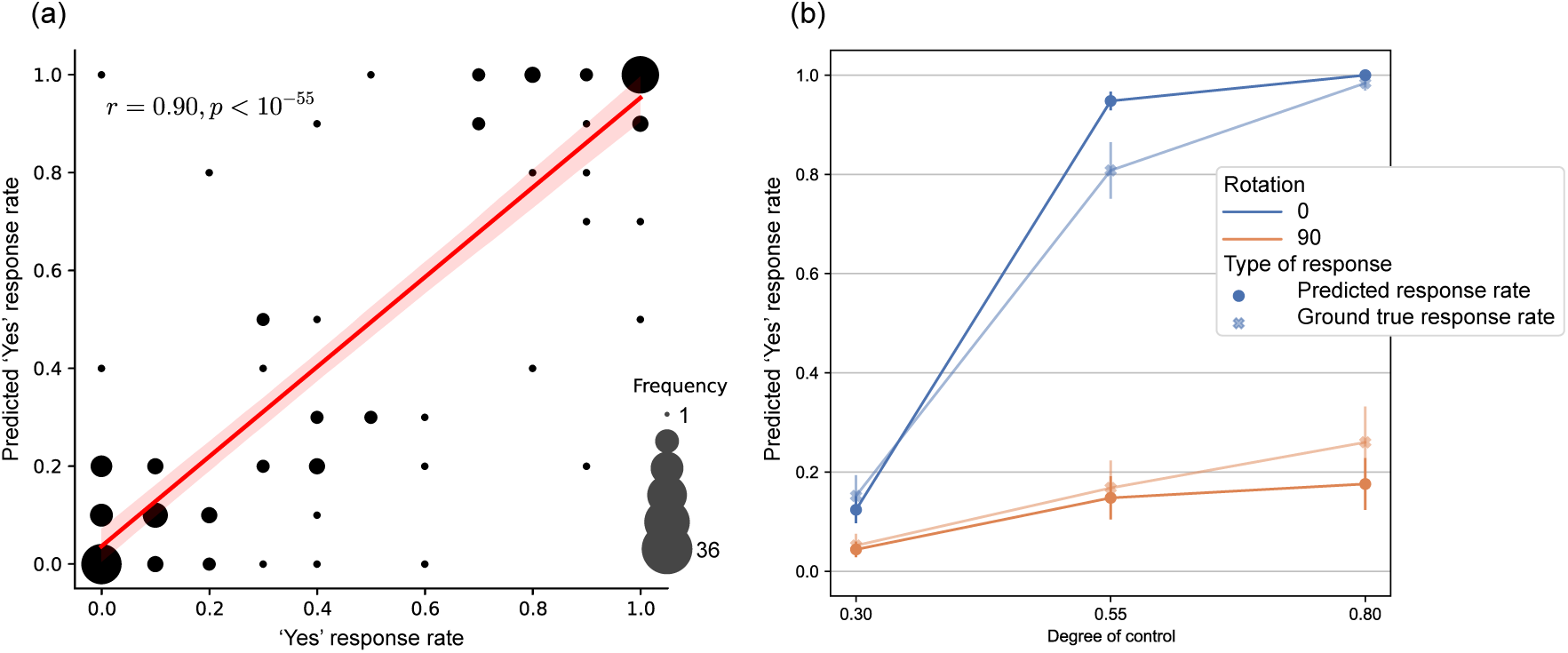
(a) The correlation between predicted response rates and actual ’Yes’ response rates cross conditions and participants in the control judgment task.(b) The group level response aggregating the predictive accuracy and the ground truth accuracies in each condition.

**Figure S5:**
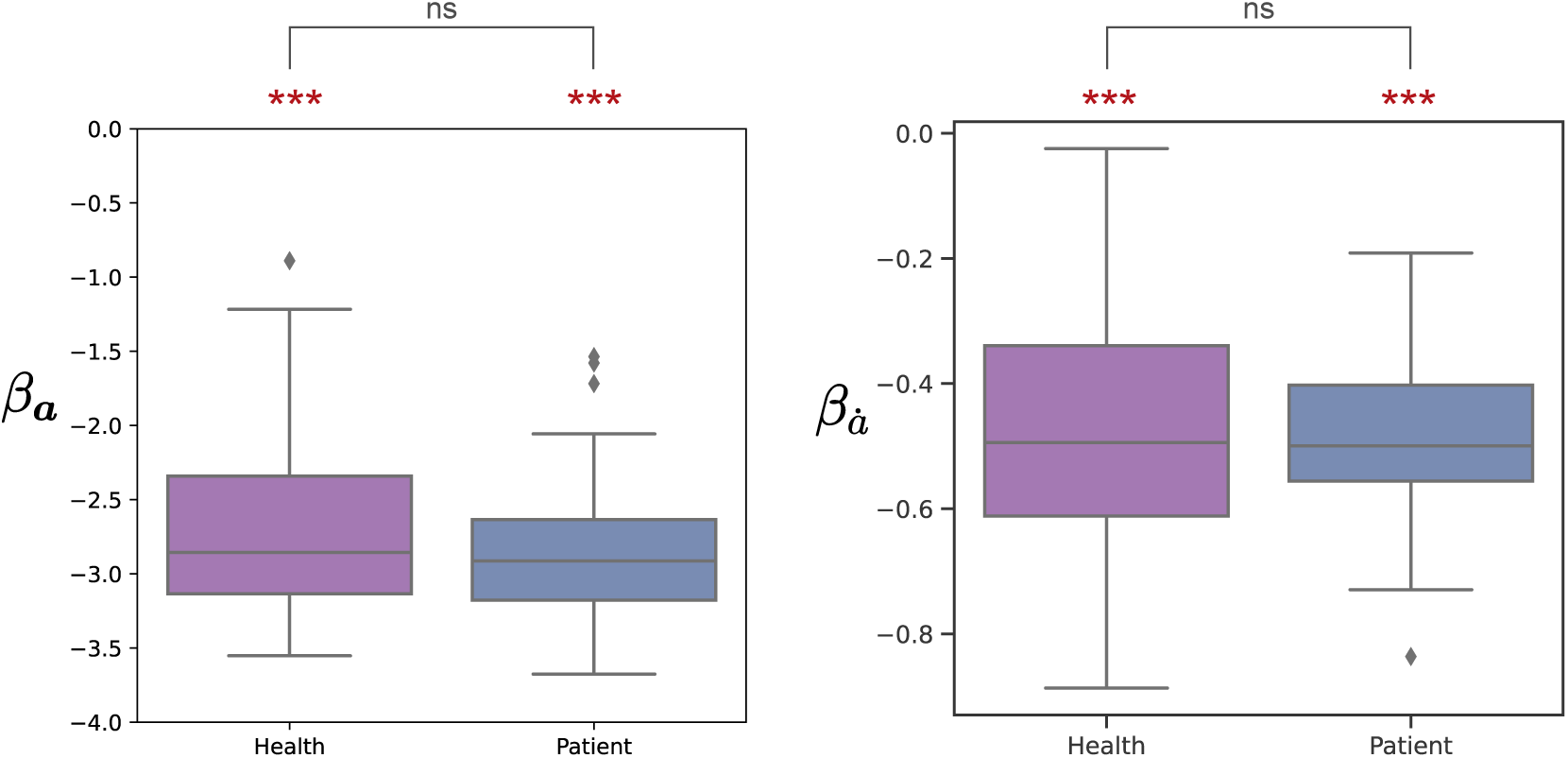
Logistic regression coefficients for action plans in the control judgment task. Presented are box plots contrasting the coefficients for the action plan distance (*β_α_*, left) and distance of action plan dynamics (*β_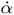_*, right) between groups: healthy subjects (depicted in purple) and schizophrenia patients (shown in blue). All coefficients are highly significantly below zero (*** *p <* 10*^−^*^10^). No significant difference was found between the two groups in either coefficient.

